# Quantitative imaging of *Caenorhabditis elegans* dauer larvae during cryptobiotic transition using optical diffraction tomography

**DOI:** 10.1101/2021.04.26.441445

**Authors:** Kyoohyun Kim, Vamshidhar R. Gade, Teymuras V. Kurzchalia, Jochen Guck

**Affiliations:** Biotechnology Center, Center for Molecular and Cellular Bioengineering, Technische Universität Dresden, 01307 Dresden, Germany; Max Planck Institute for the Science of Light & Max-Planck-Zentrum für Physik und Medizin, 91058 Erlangen, Germany; Max Planck Institute of Molecular Cell Biology and Genetics, 01307 Dresden, Germany; Institute of Biochemistry, ETH Zürich, 8093 Zürich, Switzerland

## Abstract

Upon starvation or overcrowding, the nematode *Caenorhabditis elegans* enters diapause by forming a dauer larva. This larva can further transit into an anhydrobiotic state and survive harsh desiccation. We previously identified the genetic and biochemical pathways essential for survival — but without an accompanying physical model, the mechanistic understanding of this amazing phenomenon will remain inadequate. Neither microscopic investigation of structural changes upon entry into anhydrobiosis nor the most basic quantitative characterization of material properties of living desiccated larvae, however, have been feasible, due to lack of appropriate techniques. Here, we employed optical diffraction tomography (ODT) to quantitatively assess the internal mass density distribution of living larvae in the reproductive and diapause stages. More importantly, ODT allowed for the first time physical analysis of desiccated dauer larvae: their mass density was significantly increased in the anhydrobiotic state. We also applied ODT on different mutants that are sensitive to desiccation. Remarkably, one of them displayed structural abnormalities in the anhydrobiotic stage that could not be observed either by conventional light or electron microscopy. Our advance opens a door to quantitatively assessing fine differences in material properties and structure necessary to fully understanding an organism on the verge of life and death.

## Introduction

To withstand fluctuations in environmental conditions, organisms have developed various strategies. One such strategy is entering a dormant state. The extreme form of dormancy is cryptobiosis (hidden life) when the metabolism of an organism under conditions that are not compatible with life (no food, no water or oxygen, very high or low temperatures, high osmotic pressure *etc*.) is reduced to an undetectable level. Upon encountering favorable conditions, the organism exits the cryptobiotic state and resumes metabolism and other vital activities. Prominent examples of cryptobiosis are: survival of dry bacterial or fungal spores, dormant plant seeds, or the ability of tardigrades and some nematodes to survive desiccation (1–4). Studying the molecular, structural, and material mechanisms that accompany this reversible cryptobiotic transition, is fundamental for identifying the essential differences between the living and the dead state of an organism.

In the last decade, the nematode *C. elegans* has been established as a model for studying one form of cryptobiosis — anhydrobiosis (life in the absence of water) (5). Under favorable conditions, *C. elegans* goes through the reproductive life cycle where a fertilized egg develops through larval stages from L1 − L4 to a reproductive adult. However, when encountering a harsh environment, overcrowding or food scarcity, *C. elegans* pauses the reproductive cycle and enters diapause by forming a non-feeding dauer larva (6). It has been shown that this dauer larva can survive severe desiccation (5), high osmotic pressure, or freezing (7). The dauer larvae differ from reproductive larvae in both metabolism and morphology (6, 8). They have reduced metabolic activity (oxygen consumption rate, heat production) and as a non-feeding stage, they mostly rely on internal reserves (triacylglycerols) by utilizing glyoxylate shunt to synthesize sugars (8). Morphologically, they differ from reproductive larvae by a significant reduction in volume, which is a result of a radial shrinkage during the formation of the dauer larva.

In order to survive harsh desiccation, dauer larvae first need to be exposed to a mild decrease of relative humidity (RH), a process called preconditioning. Previously, we identified genetic and biochemical pathways that are activated during the preconditioning (9) and are crucial for survival. Among these are the many-fold increase of a disaccharide trehalose and massive biosynthesis of an intrinsically disordered protein LEA-1 (5, 9). Despite of this increasing insight into genetic and biochemical details, only very little is known about the actual morphological and material changes that enable the successful survival during reversible transitions. Only some gross anatomical changes, such as a reduction of the overall volume of the worms, have been reported. The detailed structural changes that take place inside the animal have been elusive because it is notoriously difficult to reliably image the process of desiccation or a desiccated worm both with fluorescence and electron microscopy. And going beyond structure, it has not been feasible so far to quantitatively map the distribution of the material properties inside the worm, which accompany, and arguably enable, the amazing transitions between metabolically active and inactive states, between moist and dry, between alive and dead, mainly due to lack of an appropriate non-invasive technique.

As a promising solution to address this paucity, optical diffraction tomography (ODT) has recently been developed to quantitatively map the mass density distribution inside biological specimens (10, 11). By employing interferometric microscopy, ODT can determine the three-dimensional (3D) refractive index (RI) distribution of the specimen with diffraction-limited spatial resolution (∼ 100 nm). Sine RI is roughly isomorphic to electron density, ODT offers an unbiased and label-free view into the structure of living organisms. Moreover, this structure directly translates into quantitative mass density distributions, since RI and density of materials present in biological samples are linearly proportional (12, 13). While ODT has been extensively used for characterizing the mass density distribution inside individual cells (14, 15), its application on larger tissues and whole organisms has hardly been explored.

Here, we show that ODT can be employed for imaging the 3D RI distribution in living *C. elegans* larvae with clearly visible morphological features. Reconstructed RI tomograms allowed us to assess the internal mass density distribution, dry mass, and volume of larvae in the reproductive, diapause and, most importantly, in desiccated stages. The latter gave a unique opportunity to quantify the physical properties of an intact living organism in a desiccated state. We found that the mass density of *C. elegans* larvae increased upon entry into dauer diapause — due to radial volume shrinkage at constant dry mass. Further, the dauer larvae in their anhydrobiotic state exhibited very high RI values (*n* ∼ 1.5). This value is comparable to that of glass, and rarely seen in biological objects. The desiccated dauer larvae recovered their original volume in response to rehydration, but with significantly reduced dry mass (∼ 25%) and mass density. We also applied ODT to image several mutants that are sensitive to desiccation. Remarkably, one of them, *lea-1*, showed structural defects in the form of void regions with low mass density, which were not detected by other microscopy techniques. Thus, ODT is able to capture the global as well as detailed local changes of biophysical properties throughout the entire larva. We used this method to quantitatively map the material and structural changes accompanying the anhydrobiotic transition in dauer larvae. Our findings open a door to quantitatively understanding the interdependence of material properties of an organism in relation to growth, diapause and cryptobiotic states.

## Results

### Quantitative refractive index and mass density imaging of *C. elegans* larvae

We used optical diffraction tomography (ODT), employing Mach-Zehnder interferometric microscopy (Fig. 1A, see Methods) to image the spatial distribution of refractive index (RI) inside living *C. elegans* larvae. ODT reconstructed the 3D RI distribution of the specimen with diffraction-limited resolution (ca. 120 nm and 440 nm in the lateral and axial direction, respectively) from 2D quantitative phase images obtained from various incident angles. The whole-organism tomogram was enabled by stitching together RI tomograms of multiple fields of view. The mass density was directly calculated from the reconstructed RI tomograms since the RI of most biological samples, *n*_sample_, is linearly proportional to the mass density, *ρ*, as *n*_sample_ = *n*_m_ + *αρ*, where *n*_m_ is the RI of medium and *α* is the RI increment (*dn/dc*) with *α* = 0.190 mL/g for proteins and nucleic acids (16, 17).

**Figure 1.**
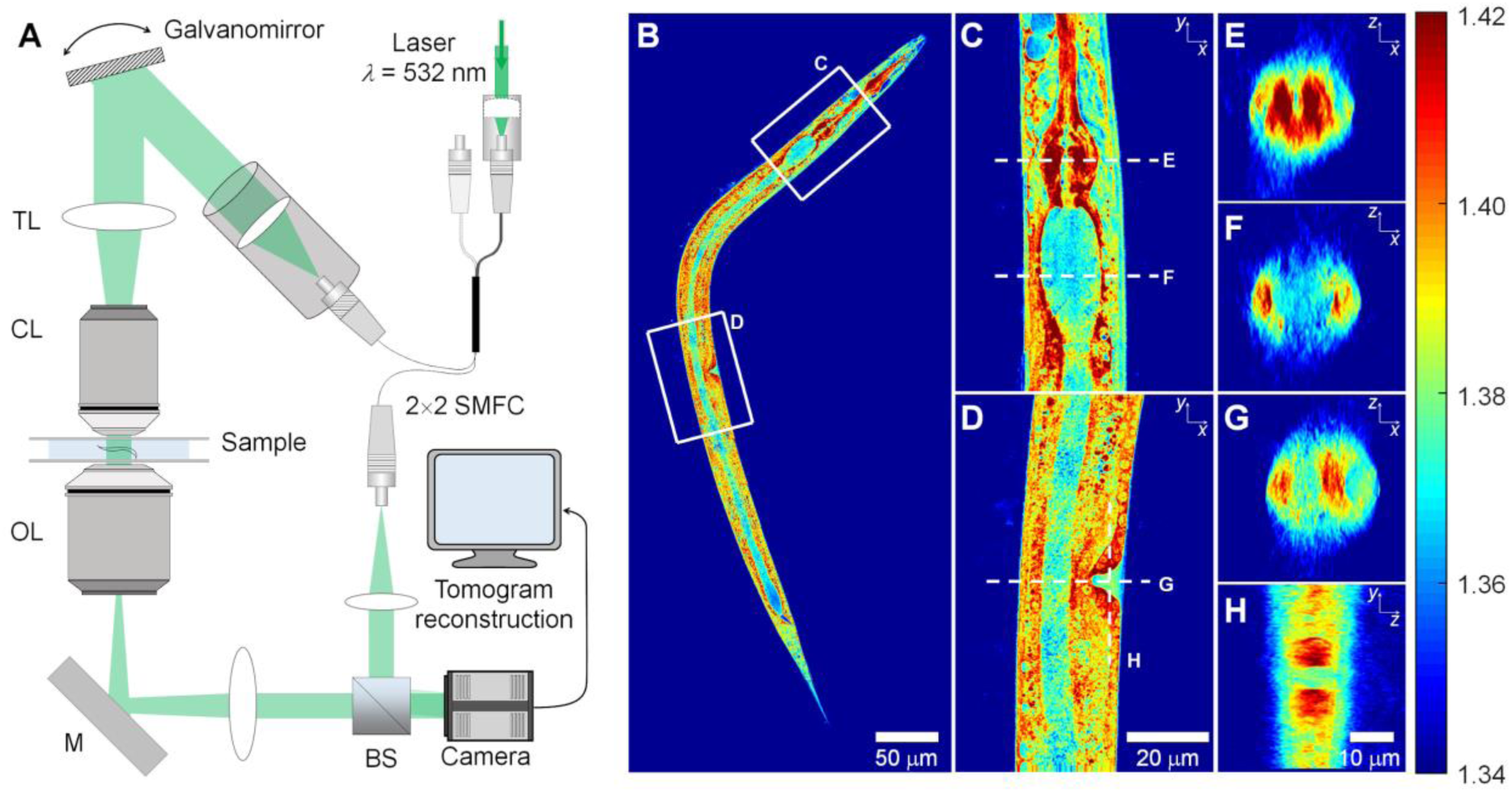
Experimental setup and representative 3D RI tomogram of a *C. elegans* larva. (A) The setup for optical diffraction tomography (ODT). SMFC: single-mode fiber coupler, TL: tube lens, CL: condenser lens, OL: objective lens, M: mirror, and BS: beam splitter. (B) Central cross-sectional slice through a 3D RI tomogram along the *x*-*y* plane of a larva at the L3 stage, and (C, D) the enlarged cross-sectional RI slices of the pharynx and vulva region indicated in (B). (E, F) Cross-sectional RI slices of (E) pharynx and (F) gut lumen along the *x*-*z* plane indicated in (C). (G, H) Cross-sectional RI slices of the vulva region along (G) the *x*-*z* plane and (H) *y*-*z* plane indicated in (D). Color scale shows RI.

Representative high-resolution images of RI tomograms of a larva at the L3 stage shown in Figure 1B – H and Supplementary Video 1 show that ODT can reveal various morphological structures in the RI contrast. Very clearly distinguishable are pharynx (with metacorpus and terminal bulb) and gut (Fig. 1C and 1E – F). Cells of the latter contain lipid droplets with very high RI, whereas the gut lumen exhibits a much lower RI. Interestingly, the pharynx is a tube formed by very tightly packed muscles, and has a similar RI value to that of lipid droplets. In addition, the vulva, having muscles, exhibits a higher RI than the surrounding tissue (Fig. 1D and 1G – H).

Next, we set out to investigate the material properties of different larval stages of *C. elegans* using ODT. For this we took advantage of the *daf-2(e1370)* strain, which enters the reproductive life cycle (L2, L3 larvae) when exposed to 15°C, but which forms dauer larvae at 25°C via an L2d intermediate (Figure 2A, B). We began with larvae at reproductive larval stages (L1, L2, and L3). The representative RI tomograms in Figure 2A clearly show detailed morphological structures in the larvae. The tomograms also show that the larvae grow in size during the reproductive cycle while maintaining a similar RI value throughout.

**Figure 2.**
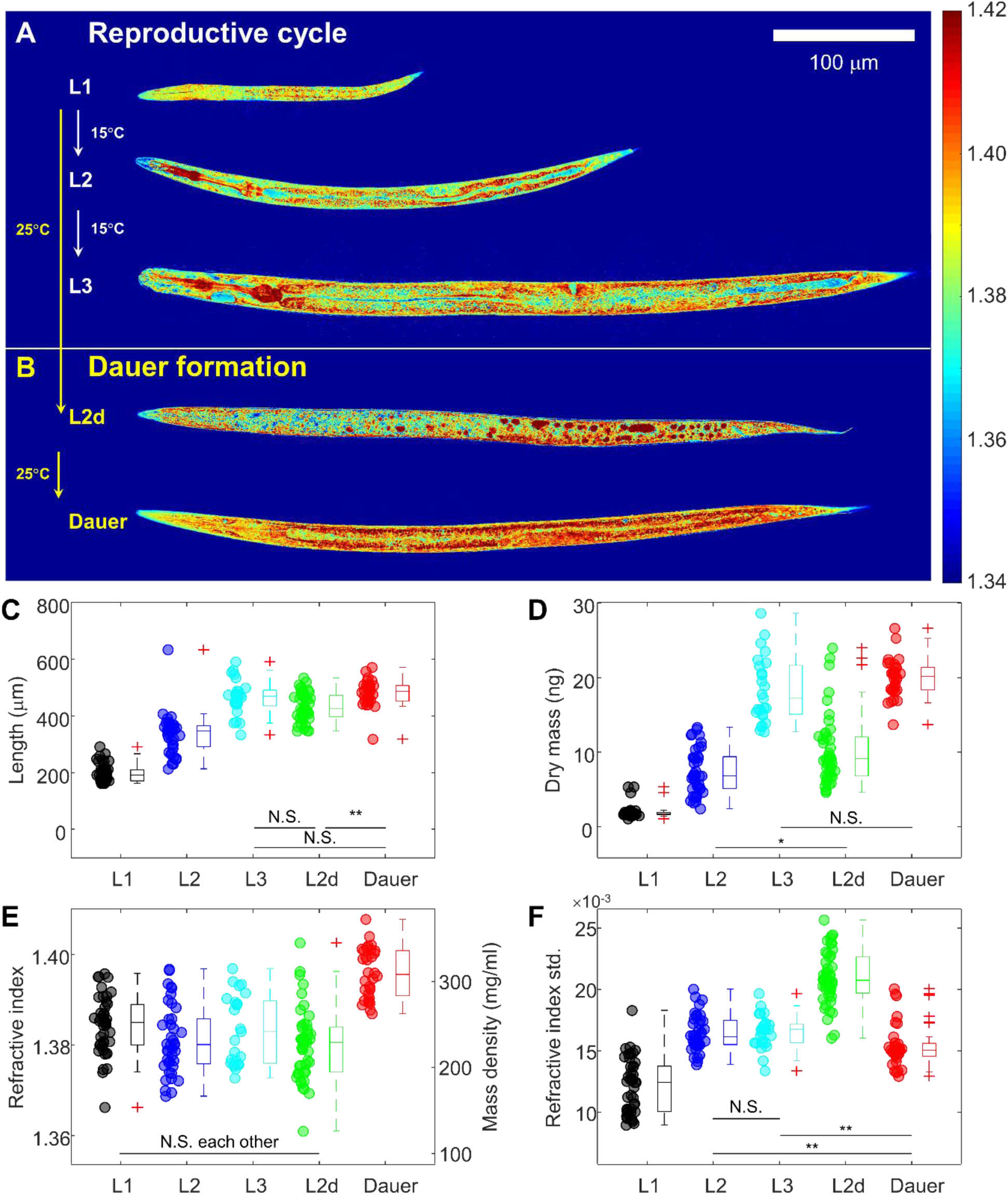
Quantitative physical analysis of *C. elegans* larvae at different larval stages. (A) Representative central cross-sectional slices through RI tomograms of *C. elegans* larvae at L1, L2, and L3 larval stages in the reproductive cycle. (B) Cross-sectional slices through RI tomograms of *C. elegans* larvae at the L2d and dauer stages. Color scale shows RI. (C – F) The length (C), dry mass (D), mean RI and mass density (E), and standard deviation of RI (F) of *C. elegans* larvae at different larval stages. The numbers of larvae measured are *N* = 43, 40, 25, 45, and 32 for L1, L2, L3, L2d, and dauer, respectively.

By quantitative analysis of the reconstructed tomograms, we characterized length, dry mass and mean RI value of the larvae (see Methods). As expected, both length and dry mass increased during growth from L1 to L3 (Fig. 2C, D). Importantly, however, the mean RI value and therefore the mean mass density of the larvae did not change significantly (1.3845 ± 0.0020, 1.3812 ± 0.0024, and 1.3835 ± 0.0032 at L1, L2, and L3 stage, respectively; corresponding to the mass density of 249.8 ± 10.4 mg/ml, 232.4 ± 12.7 mg/ml and 244.9 ± 16.6 mg/ml; Figure 2E). These measurements indicate the presence of regulatory mechanisms coordinating the biosynthesis of different classes of molecules required for growth and maintenance of metabolism and structure of cells. Moreover, the standard deviation of RI inside individual larvae at the L1 stage was as low as 0.0120 ± 0.0007 and increased to 0.0164 ± 0.0005 and 0.0165 ± 0.0006 at the L2 and L3 stages, respectively (Fig. 2F). The increasing heterogeneity of RI and mass density quantifies the developmental growth and maturation of organs such as pharynx (with metacorpus and terminal bulb) and gut.

### Dauer larvae have higher mass density than larvae in the reproductive cycle

As a next step, we investigated the RI distribution during the transition into diapause. As shown in Figure 2B and 2E, the pre-dauer larval stage called L2d larvae exhibited a similar average RI value (1.3802 ± 0.0024) to that of larvae in the reproductive cycle. However, the dauer larvae showed significantly higher RI values (1.3960 ± 0.0020). The increased RI, and mass density, of the dauer larvae might correlate with both radial shrinkage and increased accumulation of lipid droplets, known to occur during the transition to diapause. We estimated to which extent the accumulation of lipid droplets contributes to the increase in RI and mass density. For this, we correlated the RI tomograms with epi-fluorescence images of dauer larvae stained with Nile Red for lipid droplets in the same optical setup (see Methods, Supplementary Fig. 1A, B). The mean RI value of the regions containing lipid droplets was 1.4184 ± 0.0025, which is higher than non-lipid regions with 1.4081 ± 0.0010 (Supplementary Fig. 1C). In addition, the mean RI value of non-lipid regions by itself was already significantly higher than that of the L3 larvae in the reproductive cycle. Altogether our findings suggest that the increased RI in the dauer larva originates from both volume shrinkage and lipid droplet accumulation. The first increases the overall RI and mass density, whereas the latter contributes to the additional RI increase.

### Entry into anhydrobiotic state increases RI and mass density of dauer larvae dramatically

The most interesting findings were revealed in the investigation of morphological and biophysical changes in dauer larvae in a desiccated state. As previously described (5) dauer larvae need to be preconditioned (mild desiccation at 98% RH) to survive harsh desiccation (60% RH; see Figure 3A). In the process, the larvae lose up to 80% and over 95% of their body water, respectively. As shown in Figure 3B – D, after desiccation the RI of dauer larvae displayed surprisingly high RI values: after preconditioning the mean RI was 1.4955 ± 0.0043 and after harsh desiccation 1.4899 ± 0.0028, respectively (Figure 3E). The increase in RI was so high that to reduce the mismatch between the larvae and the surrounding medium, we immersed the desiccated larvae into glycerol as an imaging medium (*n* = 1.4527). The significant increase of the mean RI in the desiccated dauer larvae was mainly due to the about 2.3 to 2.4-fold decrease in volume (Figure 3F). In contrast, the dry mass of the dauer larvae decreased only slightly from 17.1 ± 0.9 ng to 16.1 ± 1.2 ng and 15.2 ± 1.3 ng (Figure 3G).

**Figure 3.**
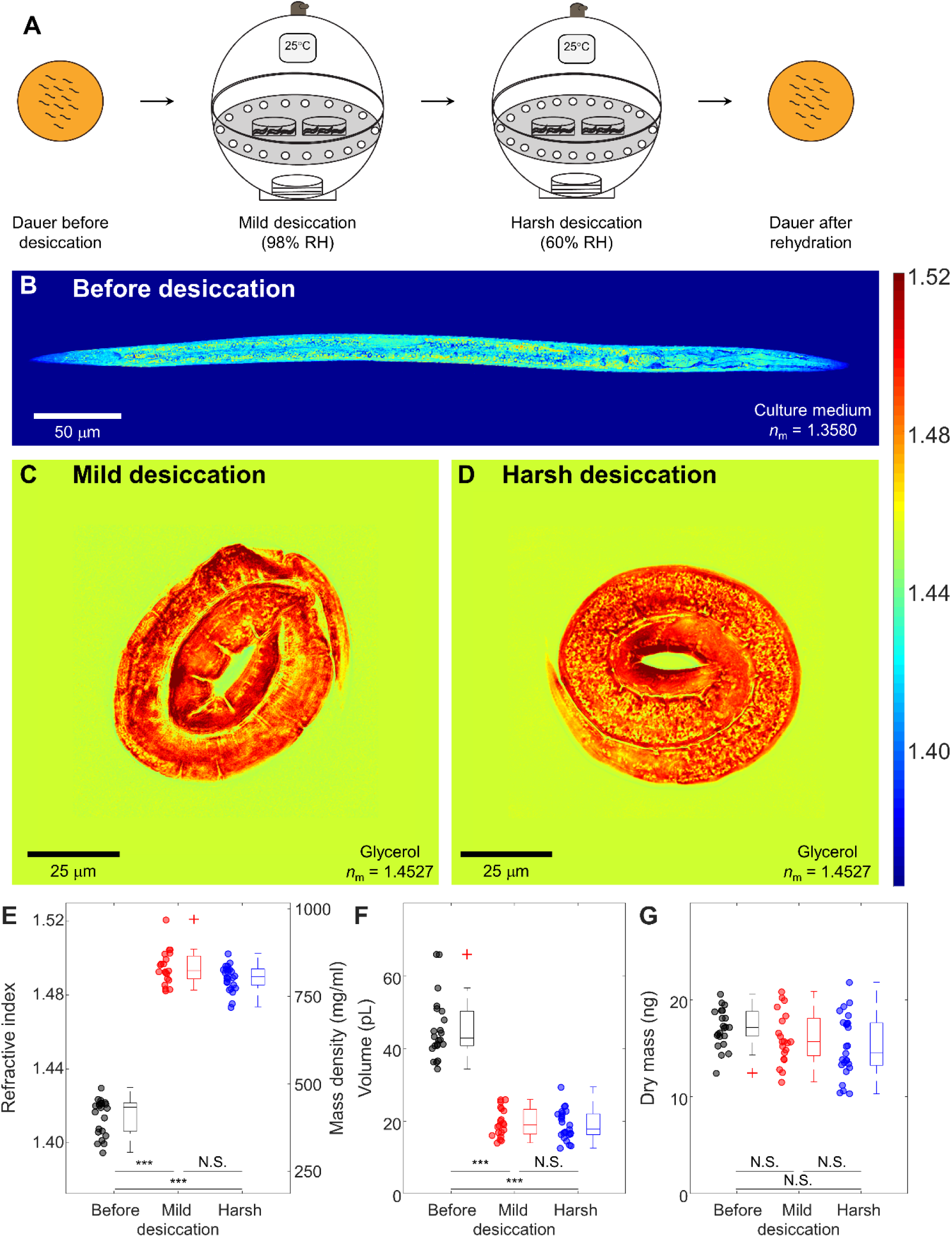
Quantitative ODT analysis of desiccated dauer larvae. (A) Schematic diagram for preparing *C. elegans* dauer larvae by consecutive mild (98% relative humidity, RH) and harsh desiccation (60% RH). (B) Central cross-sectional slice through RI tomogram of a typical *C*. *elegans* dauer larva (similar to Figure 2B, but with different RI scale). (C – D) Representative cross-sectional slices through RI tomograms of desiccated dauer larvae after (C) mild (98% RH) and (D) harsh (60% RH) desiccation. Color scale shows RI. (E – G) Mean RI (E), volume (F), and dry mass (G) of *C. elegans* dauer larvae in different desiccated stages. The numbers of desiccated larvae measured are *N* = 22, 20, and 25 for dauer, mild and harsh desiccated state, respectively.

Most cells and tissues display RI values ranging from 1.35 to 1.39 (18). Reported exceptions are diatoms whose cell walls are made of silica glass that has a high RI value of 1.46 and the basalia spicules of some glass sponges, which can reach RI values of 1.48 in their core (19, 20). Remarkably, desiccated dauer larvae had an average RI of almost 1.50 (Figure 3D), with some internal regions reaching 1.52 (Figure 3C, D), considerably higher than anything reported for other biological specimens. Thus, desiccated larvae have optical properties similar to that of glass, and quite unlike living matter. This, together with the transition of the cytoplasm from a liquid to a solid-like state (due to loss of 95% of body water), seem to be physical measures of the fact that desiccated *C. elegans* larvae are indistinguishable from dead objects.

Upon rehydration with water, the desiccated dauer larvae can revive within a few hours (5) and can develop further into reproductive adults under optimum conditions. As desiccation followed by rehydration induces breakdown of several biomolecules (*e*.*g*., triacylglycerols and trehalose) (5, 8), we hypothesized that the rehydrated dauer larvae might have different RI values and dry mass in comparison to the dauer larvae before desiccation. As shown in Figure 4A, B for typical RI tomograms and Figure 4C for quantitative results, the rehydrated dauer larvae had a mean RI value of 1.3971 ± 0.0022, which was significantly lower than that of the dauer larvae before desiccation with 1.4150 ± 0.0013. The rehydrated dauer larvae recovered the volume of dauers before desiccation (53.0 ± 2.2 pL and 55.5 ± 3.3 pL, respectively; Figure 4D). Interestingly, the dry mass of the dauer larvae decreased by almost 25% from 20.4 ± 0.6 ng before desiccation to 16.1 ± 0.8 ng after rehydration (Figure 4E). This result quantitatively confirms our previous findings that the dauer larvae consume significant amounts of triacylglycerols and trehalose during desiccation and rehydration — they metabolize a quarter of their own internal contents in the process.

**Figure 4.**
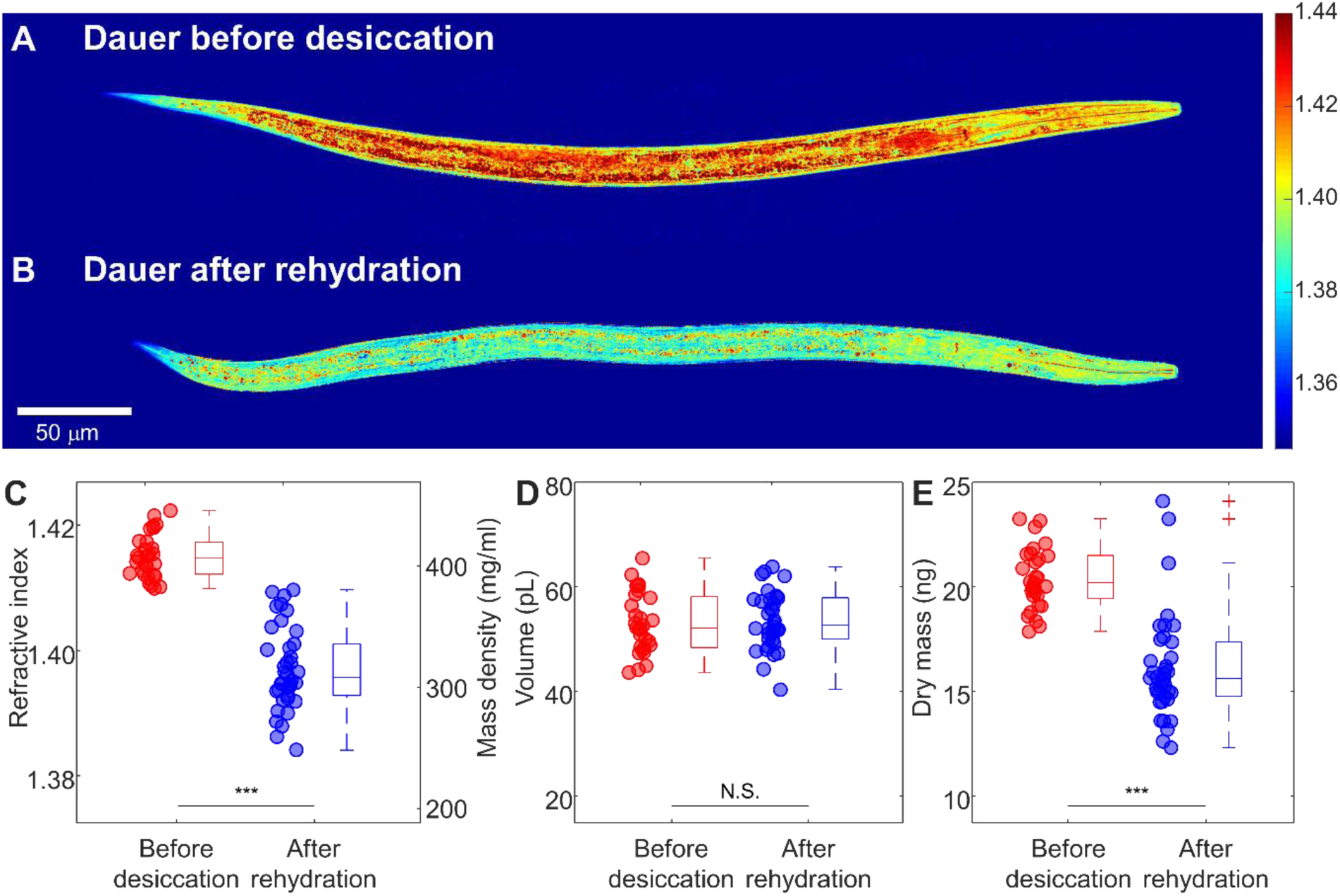
Quantitative ODT analysis of dauer larvae before desiccation and after rehydration. (A, B) Central cross-sectional slices of RI tomograms of typical *C. elegan*s dauer larvae (A) before desiccation and (B) after rehydration. Color scale shows RI. (C – E) Mean RI and mass density (D), volume (E), and dry mass (F) of *C. elegans* dauer larvae before desiccation and after rehydration. The numbers of dauer larvae measured are *N* = 29 and 38, respectively.

### ODT reveals structural differences in desiccation-sensitive mutants

Previous studies have shown that *C. elegans* dauer larvae activate several biochemical pathways to survive harsh desiccation (9). One of them leads to massive biosynthesis of the disaccharide trehalose, which is involved in protecting phospholipid bilayers against water-induced damage during rehydration (5). Another essential factor for survival is biosynthesis of an intrinsically disordered protein LEA-1 (late embryogenesis abundant) (7, 9). Deletion mutants with the biosynthetic pathways of trehalose (double mutant lacking biosynthetic enzymes TPS-1 and TPS-2, *daf-2;ΔΔtps*) or of LEA-1 (*daf-2;lea-1*) abolished are non-viable after rehydration (7). Thus, we investigated how deficiency in trehalose and LEA-1 in these mutants influences the physical characteristics of desiccated dauer larvae. Is there a correlation between the latter and the viability of larvae?

As shown in Figure 5 and Supplementary Video 2 – 4 for the representative ODTs and Supplementary Fig. 2A for the quantitative analysis, the trehalose and LEA-1 deletions did not affect overall mean RI value and mass density significantly — neither before nor after mild and harsh desiccation. Only the dauer larvae of the *lea-1* deletion mutant had a slightly lower mean RI value than wild type dauer larvae. Not surprising for mutants that cannot produce trehalose, the volume and dry mass of the dauer larvae before and during desiccation were lower than that of control dauer larvae in the same conditions (Figure 5D – F and Supplementary Fig. 2B, C). The result is consistent with previous studies showing that trehalose is produced largely during the preconditioning (5) However, since volume and mass changes in the trehalose mutants scaled proportionally, the overall densities of wild type and both mutant larvae were similar in the anhydrobiotic state.

**Figure 5.**
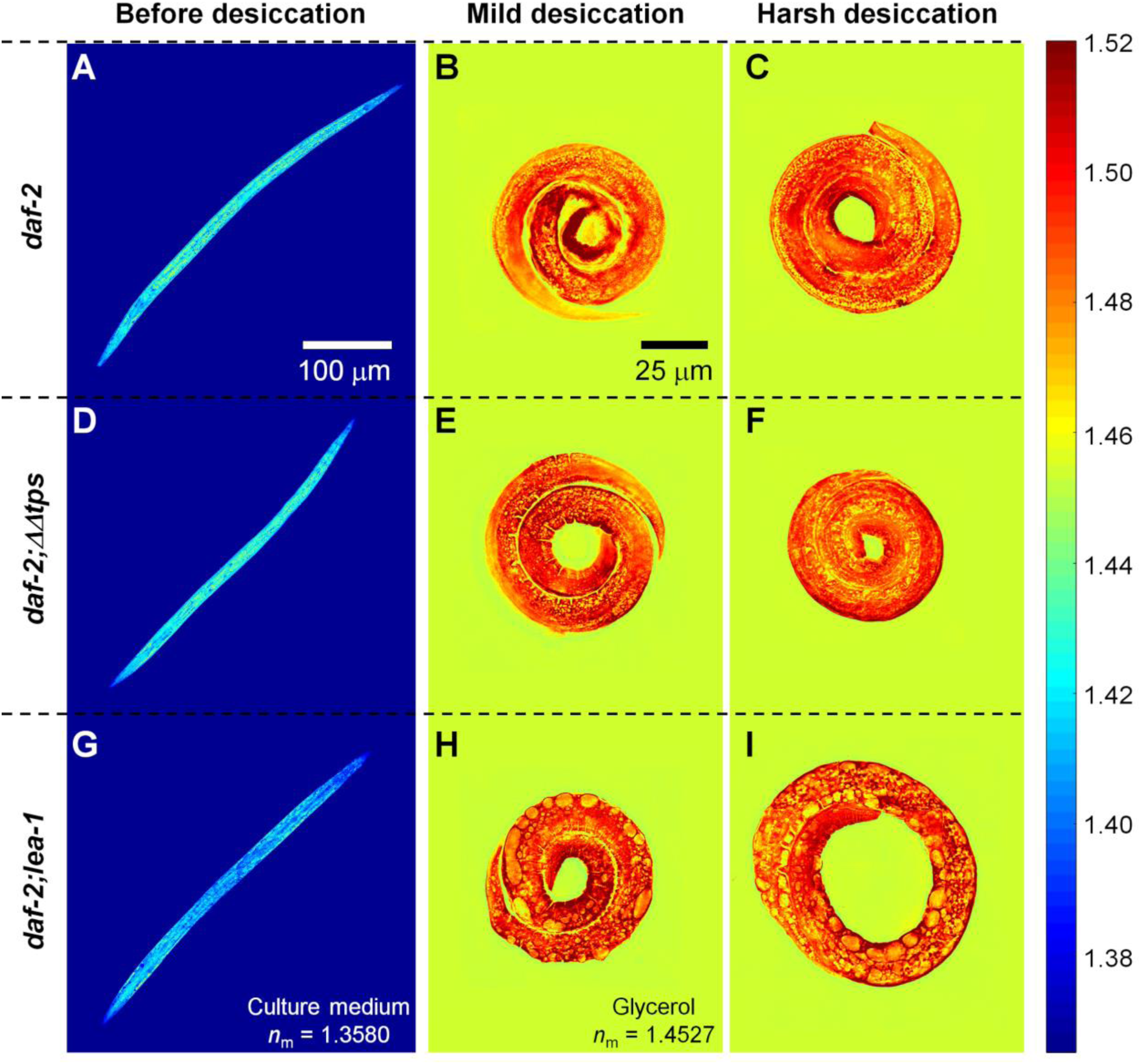
ODT analysis of desiccated dauer larvae with different genetic mutations. Typical central cross-sectional slices through RI tomograms of *C. elegans* dauer larvae of (A – C) controls, (D – F) trehalose deletion mutants *daf-2;ΔΔtps*, and (G – I) *lea-1* deletion mutants *daf-2;lea-1*, respectively. (A, D, G) represent dauer larvae before, (B, E, H) after mild (98% RH), and (C, F, I) after harsh desiccation (60% RH). Color scale shows RI.

Even though the *lea-1* mutants were inconspicuous in their overall physical properties, they displayed very interesting internal structural anomalies. As seen in Figs. 5H-I, the RI tomograms of desiccated dauer larvae of the *lea*-1 deletion mutant displayed distinct void regions with significantly lower RI value. By visual inspection, the void regions corresponded to mid gut regions anatomically. This finding based on ODT measurements quantifies morphological and physicochemical differences in desiccated samples that are hardly, or not at all revealed by light or electron microscopy (EM). As seen in Supplementary Fig. 3, bright field microscopy did not display any difference in contrast to wild type. Differential interference contrast (DIC) microscopy did show circular structures that might correspond to the void volumes detected by ODT. But without the 3D imaging and quantification capabilities of ODT, one cannot determine these differences in local material properties with certainty. Further, imaging with ODT has considerable advantages to EM stemming from the fact that it can image specimens in their native state and without any preparative steps, as in EM, that are necessarily destructive and also bear the danger of creating artifacts or eliminating differences. It must be said that EM principally does not allow the analysis of a desiccated worm as it can only be applied to samples after rehydration. The electron micrographs of the dauer larvae of *lea-1* deletion mutant after harsh desiccation, and subsequently necessary rehydration, show that the annular morphology of the desiccated larvae is distorted (Supplementary Fig. 4). This observation can, however, only indirectly indicate possible structural changes of *lea-1* dauer larvae in the desiccated state.

The measured RI tomograms were further analyzed quantitatively to investigate the mass density difference in void regions. The void regions were segmented from the RI tomograms by applying the Otsu method (21), and the mean RI value of the void regions and peripheral regions was quantified. The periphery of the void regions was segmented by dilating the binary masks for the void regions by 5 μm. The mean RI value of the void regions of the dauer larvae in mild and harsh desiccation conditions was 1.4686 ± 0.0004 and 1.4686 ± 0.0006, respectively, which were significantly lower than the peripheral regions with 1.4962 ± 0.0006 and 1.4972 ± 0.0008 (Supplementary Fig. 5). The relative RI difference suggests that the void region is 18% less dense than peripheral regions. It is remarkable that ODT can detect and quantitate such fine structural differences in an organism. Altogether our ODT-based findings shed light onto the correlation between material and structural properties of an organism and its survival ability in extreme environments.

## Discussion

In this study, we employed ODT to reconstruct the 3D RI distribution of *C. elegans* larvae in reproductive stages and dauer diapause. From these RI tomograms, the physical properties including mass density, volume, and dry mass were quantitatively analyzed. So far, ODT has mostly been applied for single-cell analysis mainly due to the limited field-of-view (14, 15), and few studies briefly visualized the RI tomograms of *C. elegans* (22, 23). This study is the first quantitative ODT analysis of an entire organism.

We used ODT to provide the biophysical and structural properties of live dauer larvae, which have not been charted by conventional microscopic techniques. Previously, EM had been used to reveal the fine structures of dauer larvae and their density differences under harsh desiccation conditions with its typical very high spatial resolution (5, 24). However, the sample preparation for desiccated dauer larvae requires rehydrating the larvae for a brief time before further processing. Thus, EM images we obtain do not depict larvae in desiccated state but rather changes that occur after first desiccation and then subsequent rehydration. ODT might not provide the same spatial resolution as EM, but it can quantitatively assess structure and physical properties of intact dauer larvae without any sample preparation steps. In principle, ODT can even trace the changes within the same larvae during development and the desiccation process. As another relevant light microscopy, fluorescence microscopy provides molecularly specific localization of fluorescently stained proteins and organelles in live dauer larvae. However, fluorescence imaging is susceptible to phototoxicity and can only visualize molecules labeled selectively. It is not suitable to quantify physical properties and their changes. In contrast, ODT allows revealing the mass density of overall unbiased substances in the entire organism. Since *C. elegans* larvae can be considered optically transparent, at least up to the L3 stage in the reproductive life cycle (25), the first-order Rytov approximation is still valid for reconstructing RI tomograms. To extend analysis to even larger and optically denser specimens, various computational algorithms for tomogram reconstruction have recently been developed to take into account multiple photon scattering in the sample (23, 26). Employing such algorithms, we can extend our studies in the future also to L4 larvae and adult *C. elegans* worms. From reconstructed RI tomograms, we found that the *C. elegans* larvae in the reproductive cycle maintain a constant mass density during development, while entry to dauer diapause increases the mass density significantly. From the correlation between the RI tomograms and epi-fluorescence images of Nile Red-stained lipid droplets, we conclude that the increased mass density in dauer larvae is presumably due to this lipid droplet accumulation, in addition to the radial shrinkage they undergo during dauer formation. This hypothesis can be tested by analyzing whether mutant strains that do not undergo proper radial shrinkage (24, 27) or that deplete lipid droplets rapidly (28) exhibit a similar increase in mass density.

Biophysical properties of organisms that can survive extreme environments have until now only sporadically been studied. One remarkable finding of our study is that dauer larvae in their anhydrobiotic state (where most of the body water is lost) have a very high RI value reaching *n* ∼ 1.5. This RI value is usually not found in biological specimens and is comparable to that of glass (Supplementary Fig. 6). While this finding needs further investigation into the biological relevance of why and how desiccated dauer larvae acquire such high RI and mass density, one can speculate that the increased mass density also corresponds to altered mechanical properties. A recent study showed that crowding above the critical mass density (∼ 340 mg/ml) induces glass-forming behavior of the cytoplasm in cells with a resulting increased viscosity (29). This mass density value corresponds to an RI value of about 1.4, which ranges between that of dauer larvae and desiccated dauer larvae as found in our study. Moreover, it has been shown that the viscoelastic property of tardigrades, one of the other species that can enter an anhydrobiotic state, becomes glass-like in desiccation (30). Therefore, the dramatic increase in RI and mass density may reflect a glass transition of the cytoplasm during desiccation. It is conceivable that such a transition contributes to an increased mechanical stability in the desiccated state of the organism. This hypothesis needs further validation by direct measurements of mechanical properties of desiccated dauer larvae. Conventional, contact-based techniques for mechanical phenotyping of biological samples with (sub-)cellular resolution, such as atomic force microscopy-enabled nanoindentation, necessitate a destruction and slicing of the sample in order to gain access to internal material properties (31). This approach would be inappropriate to obtain reliable mechanical information about the process of desiccation. However, recently, non-invasive microscopic techniques to probe the mechanical properties of biological samples directly inside living biological samples have emerged, including Brillouin microscopy (32–35) and time-lapse quantitative phase microscopy (36, 37). Combining ODT with such microscopic techniques can in the future provide decisive information on the detailed nature of the material transitions during dauer formation and desiccation of *C. elegans* larvae.

Our measurements on rehydrated dauer larvae revealed that the dry mass is significantly decreased (∼ 25%) in rehydrated larvae while their volume remains constant. It is worth noting that ODT provides absolute and unbiased quantification of how much material the larvae consume during the rehydration process. In accordance with our previous results, we found that the degradation of essential biomolecules (triacylglycerols, trehalose) upon desiccation and rehydration (5, 8) manifest on the dry mass content of the dauer larvae. The significant increase and decrease of mass density during harsh desiccation and rehydration without damage draw our attention to the connection between biochemical pathways and material properties of the larvae in such dramatic transitions. Hence, we quantitatively characterized the physical differences of deletion mutants that do not survive desiccation. To the best of our knowledge, this is the first study reporting the overall changes in the RI and mass density distributions of an entire multicellular organism with genetic mutations. The mean RI value of the deletion mutants remained the same as the wild type desiccated larvae. However, we observed a decreased dry mass and void regions with low mass density in deletion mutants. The trehalose deletion mutant (*daf-2;ΔΔtps*) displayed a decreased dry mass, which is in accordance with our previous observation that trehalose levels are normally accumulated during desiccation (5). The *lea-1* deletion mutant larvae showed distinct structural differences in the RI distribution during desiccation, as they exhibited void regions with significantly lower RI value in the RI tomograms. The molecular mechanism of how LEA-1 confers desiccation tolerance to dauer larvae remains elusive. Several *in vitro* studies have indicated that LEA-1 is involved in the prevention of protein aggregation during desiccation (38, 39). In combination with our electron microscopy results, the structural defects in the RI tomograms of *lea-1* deletion mutant indicate that LEA-1 might maintain the functionality of cytosolic proteins which further assist in the maintenance of the annular morphology of the desiccated larvae (Supplementary Fig. 4). Further correlative investigations with fluorescently tagged LEA-1 in wild type larvae and corresponding regions in *lea-1* deletion mutant should lead towards the precise mechanism.

To conclude, we utilized ODT to quantitatively investigate the physical and structural changes in a living *C. elegans* larvae during dauer formation and upon desiccation. We revealed that the RI of dauer larvae is higher than that of larvae in the reproductive cycle, and becomes even as high as the RI of glass (*n* ∼ 1.5) in the desiccated state. Moreover, dauer larvae of the deletion mutants of trehalose and LEA-1 exhibited distinct morphological changes in the desiccation condition, which may affect the survival in such harsh environments. The biological relevance of a higher mass density of the larvae during dauer formation and upon desiccation requires further investigation. However, the physical understanding and corresponding quantitative modeling of cryptobiotic transitions in *C. elegans* larvae can now be based on actual physical parameters determined by methods such as ODT. As such, our study paves the way to a more complete understanding of the underlying mechanisms to sustain the integrity of nematodes, and ultimately other organisms, in transitions between life and death.

## Methods

### Materials, *C. elegans* strains and growth conditions

The Caenorhabditis Genetic Centre (CGC) provided the *C. elegans* strain *daf-2(e1370)* and the *E. coli* strain NA22. The compound mutant strains of *daf-2(e1370)III;lea-1(tag1676)V, tps-2(ok526)II; daf-2(e1370)III; tps-1(ok373)X(daf-2;ddtps)* were generated during our previous studies (5, 7).

*daf-2(e1370)* eggs were incubated in 1X M9 buffer for a few hours at room temperature at shaking to obtain synchronized hatched L1 larvae. These L1 larvae were plated on NGM agar plates with *E. coli* NA22. Half of the plates were incubated at 15°C and the rest at 25°C for reproductive and dauer larvae formation, respectively. Larval stages were monitored, visually confirmed for respective stages, and collected from the plate.

### Desiccation of *C. elegans* dauer larvae

Larvae at various stages were collected in water and washed twice with water to remove any debris. For preparing preconditioned and desiccated larvae, a dauer suspension of 5 μl was pipetted onto a coverslip (VWR International) and exposed to 98% RH (relative humidity) for 4 days and 60% RH for 1 day subsequently. For imaging desiccated dauer larvae, the dauer larvae were immersed in glycerol (*n* = 1.4527) in order to reduce the RI difference between dauer larvae and the surrounding medium. The RI of the medium was measured using an Abbe refractometer (2WAJ, Arcarda GmbH). For imaging rehydrated larvae after desiccation, the dauer larvae were rehydrated for 2 hours with water and then anesthetized with levamisole (Sigma) prior to imaging.

### Optical setup for optical diffraction tomography

The three-dimensional (3D) refractive index (RI) distribution of *C. elegans* larvae was determined using optical diffraction tomography (ODT). The optical setup was described previously (40). Briefly, ODT employs Mach-Zehnder interferometry to measure multiple complex optical fields from various incident angles (Figure 1C). A laser beam (λ = 532 nm, frequency-doubled Nd-YAG laser, Torus, Laser Quantum Inc.) was coupled into an optical fiber and divided into two paths using a 2 × 2 single-mode fiber-optic coupler (TW560R2F2, Thorlabs). One beam was used as a reference beam and the other beam passed through a tube lens (*f* = 175 mm) and a water-dipping objective lens (NA = 1.0, 40×, Carl Zeiss AG) to illuminate the sample on the stage of a home-built inverted microscope. The beam diffracted by the sample was collected with a high numerical-aperture objective lens (NA = 1.2, 63×, water immersion, Carl Zeiss AG) and a tube lens (*f* = 200 mm). To reconstruct a 3D RI tomogram of the sample, the sample was illuminated from 150 different incident angles scanned by a dual-axis galvano-mirror (GVS012/M, Thorlabs Inc.) located in the conjugate plane of the sample. The diffracted beam interfered with the reference beam at an image plane, and generated a spatially modulated hologram, which was recorded with a CCD camera (FL3-U3-13Y3M-C, FLIR Systems, Inc.). The total magnification of the setup was 57×, and the field-of-view (FOV) of the camera covers 86.2 μm × 86.2μm.

### Tomogram reconstruction and quantitative analysis

The complex optical fields of light scattered by the samples were retrieved from the recorded holograms by applying a Fourier transform-based field retrieval algorithm (41). To measure the 3D RI tomograms of whole larvae and desiccated dauers, whose size is much larger than the FOV, segmented complex optical fields of the samples were measured and digitally stitched by a custom-made MATLAB script. The 3D RI distribution of the samples was reconstructed from the retrieved complex optical fields via the Fourier diffraction theorem, employing the first-order Rytov approximation (10, 42). A more detailed description of tomogram reconstruction can be found elsewhere (43).

On the reconstructed tomograms, Otsu’s thresholding method (21) was used to segment the region occupied by the larvae from the background, and quantitative analysis was performed to calculate mean RI value, dry mass, volume, and the standard deviation of RI in the individual larvae. The mass density of the larvae was directly calculated from the mean RI value, since the RI value in biological samples, *n*(*x,y,z*), is linearly proportional to the mass density of the material, *ρ*(*x,y,z*), as *n*(*x,y,z*) = *n*_m_ + *αρ*(*x,y,z*), where *n*_m_ is the RI value of the surrounding medium and *α* is the RI increment (*dn/dc*) with *α* = 0.190 mL/g for proteins and nucleic acids (16, 17). The RI of the medium was measured using an Abbe refractometer (2WAJ, Arcarda GmbH). The volume of the larvae was extracted by counting the number of voxels in the segmented region and the dry mass of the larvae was calculated by integrating the mass density inside the segmented region. All tomogram acquisition and data analysis were performed using custom-written MATLAB scripts (R2018b, MathWorks, Inc.), which are available upon request. Tomogram rendering was performed by an open-source software (tomviz 1.9.0, https://tomviz.org/). The RI tomograms of all larvae presented in the current study are available from figshare under the following link: https://doi.org/10.6084/m9.figshare.14483331.

### Electron microscopy of desiccated dauer larvae

*daf-2* and *daf-2; lea-1* dauers that were non-preconditioned, preconditioned (98%RH) and desiccated (60%RH) were rehydrated with distilled water for 20 min after which water was soaked off and bovine serum albumin solution was added. These dauer samples were then transferred to carriers of 3 mm diameter and 0.1 mm depth and rapidly frozen in a high-pressure freezing machine (Leica, EM ICE). For automated freeze substitution, frozen samples from the above step were transferred into vials containing a special freeze substitution cocktail (Acetone, 1% Osmium tetroxide, 0.1% Uranyl acetate) by increasing the temperature to 4.5°C. After thawing, samples were rinsed with acetone to remove any freeze substitution cocktail. Then the samples were infiltrated with Epon LX112 resin: Acetone solution (1:2, 1:1, 2:3) for 1.15 h, 1.30 h, 2 h respectively. Finally, they were left in pure resin overnight and then for 4 h. After polymerization and embedding, sections of 70 nm thickness were taken with an ultramicrotome (Leica, UCT) and these sections were incubated in 1% Uranyl acetate in 70% methanol for ten minutes, followed by several washes in 70% methanol, 50% methanol, 30% methanol and finally with distilled water. Sections were further incubated in Lead citrate for 5 minutes, followed by washes with distilled water. Sample sections were analysed with an electron microscope (Tecnai12, Philips) and images were acquired with TVIPS camera (Tietz).

### Lipid droplet staining and imaging

*daf-2(e1370)* eggs were plated on NGM agar plates with *E. coli* NA22 mixed with Nile Red (Thermo scientific, 200 μg/ml). These plates were incubated at 25°C for dauer formation. After three days, dauer formation was visually confirmed. Dauer larvae were collected from the plates, washed thrice with water at 1500 g for 1 min to remove any debris and excess dye adhering to the larvae. The dauer larvae were anaesthetized with levamisole (Sigma) and imaged.

Fluorescence emission intensity of Nile red-stained dauer larvae was measured by epi - fluorescence microscopy combined in the same optical setup as ODT. The detailed configuration was described elsewhere (44). The incoherent light from a halogen lamp (DC-950, Dolan-Jenner Industries Inc.) was passed through a bandpass filter (bandwidth *λ* = 545 ± 25 nm, Carl Zeiss AG), and coupled into the same beam path in the ODT using a three-channel dichroic mirror (FF409/493/596-Di01-25×36, Semrock Inc.). The fluorescence emission signal from Nile Red in lipid droplets was collected by the same objective lens and acquired using the ODT camera. A bandpass filter (bandwidth *λ* = 605 ± 70 nm, Carl Zeiss AG) was placed in front of the camera to suppress the excitation beam. The lipid regions were segmented from measured epi-fluorescence images by applying Otsu’s thresholding method and correlated with the cross-sectional slices of RI tomograms, from which the mean RI values of lipid and non-lipid regions were calculated.

### Statistical Analysis

Measured quantities were reported as mean ± standard error of mean (SEM) throughout. Statistical significance was determined using Mann-Whitney U test. The shown asterisks indicate the statistical significance as *p < 0.01, **p < 0.001, and ***p < 0.0001, respectively.

## Acknowledgements

The authors acknowledge financial support from the Volkswagen Foundation (Life? research grant 92847). We thank Vasily Zaburdaev, Simon Alberti, Gheorghe Cojoc, Raimund Schlüßler, Titus M. Franzmann, Hui-Shun Kuan, Shada Abuhattum, and Anne Eßlinger for helpful discussions. We thank Daniela Vorkel from the electron microscopy facility of MPI - CBG for technical assistance.

## Author contributions

KK conducted the ODT measurements and analyzed the data; VG prepared the *C. elegans* larvae; KK and VG interpreted the ODT data; KK, VG, TK, and JG contributed to the conception and design of the study and interpretation of the results, and wrote the manuscript.

## Supplementary Figures

**Supplementary Figure 1.**
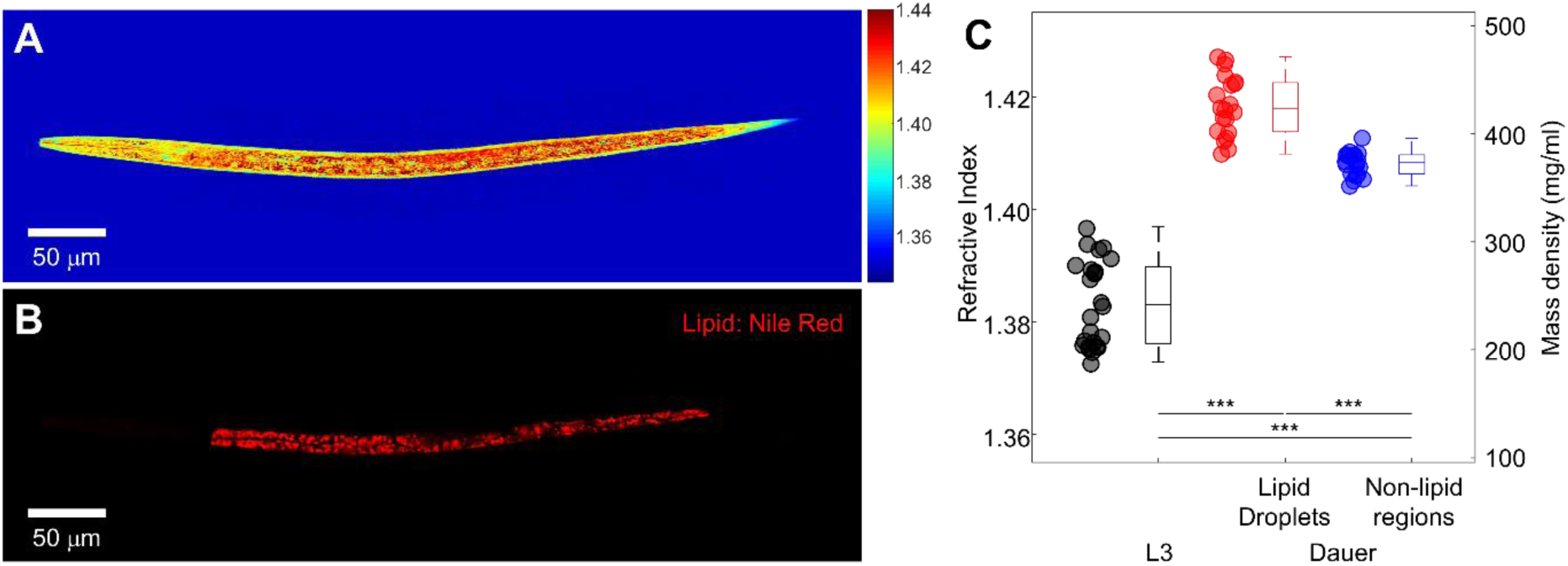
Correlation between refractive index (RI) tomograms and epi-fluorescence images of lipid droplets in dauer larvae. (A) Central cross-sectional slice through an RI tomogram of a typical dauer larva. Color scale shows RI. (B) Epi-fluorescence image of the same dauer larva in which the lipid content is stained with Nile Red. (C) Mean RI of dauer larvae at the L3 stage, as well as of lipid droplets and non-lipid regions in the Nile Red-stained dauer larvae. The numbers of L3 and dauer larvae measured are *N* = 25 and 20, respectively.

**Supplementary Figure 2.**
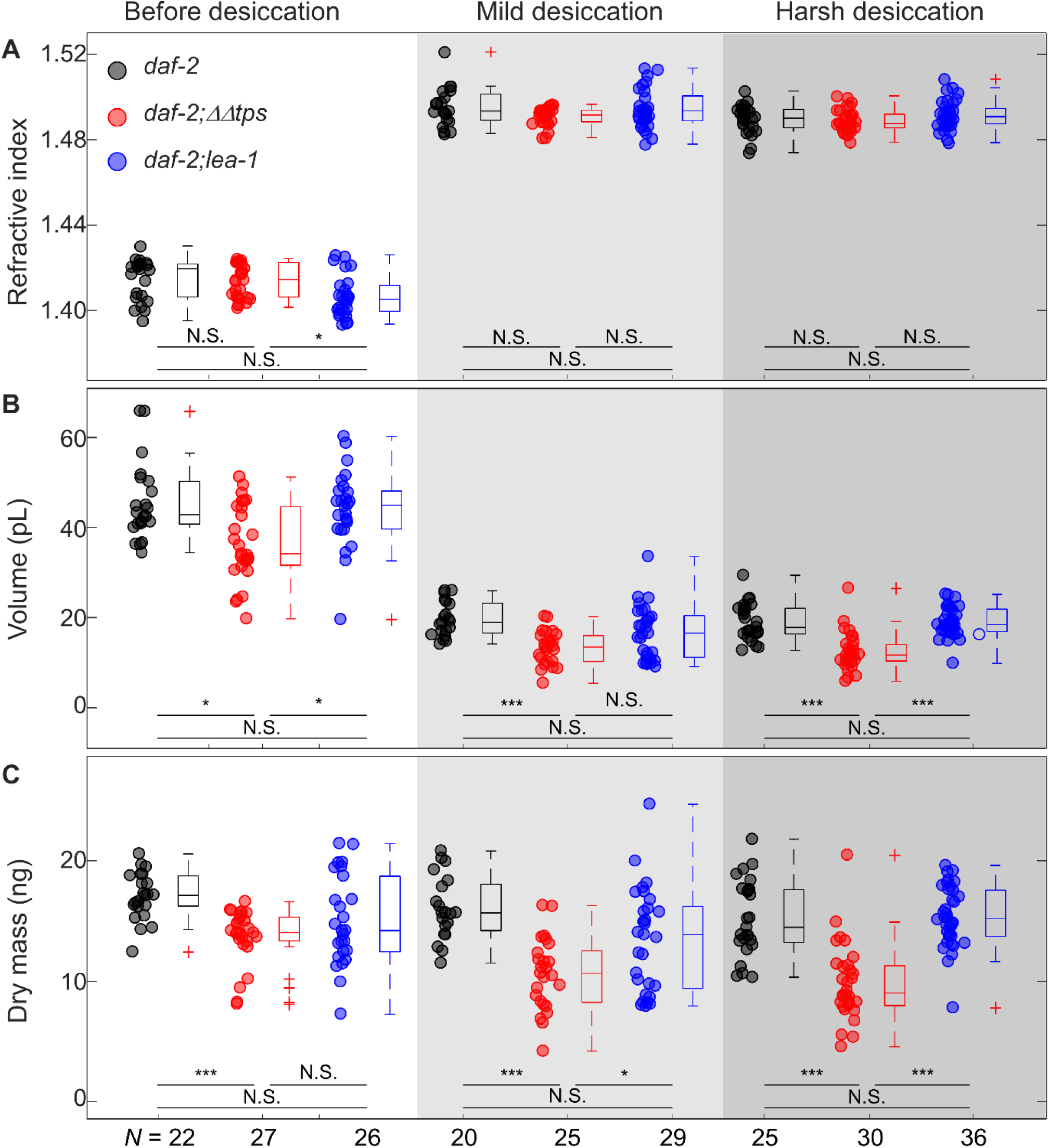
Quantitative characterization of wild type (black), trehalose deletion mutant (red), and LEA-1 deletion mutant (blue) dauer larvae in different conditions. (A) The mean refractive index, (B) volume, and (C) dry mass of the dauer larvae.

**Supplementary Figure 3.**
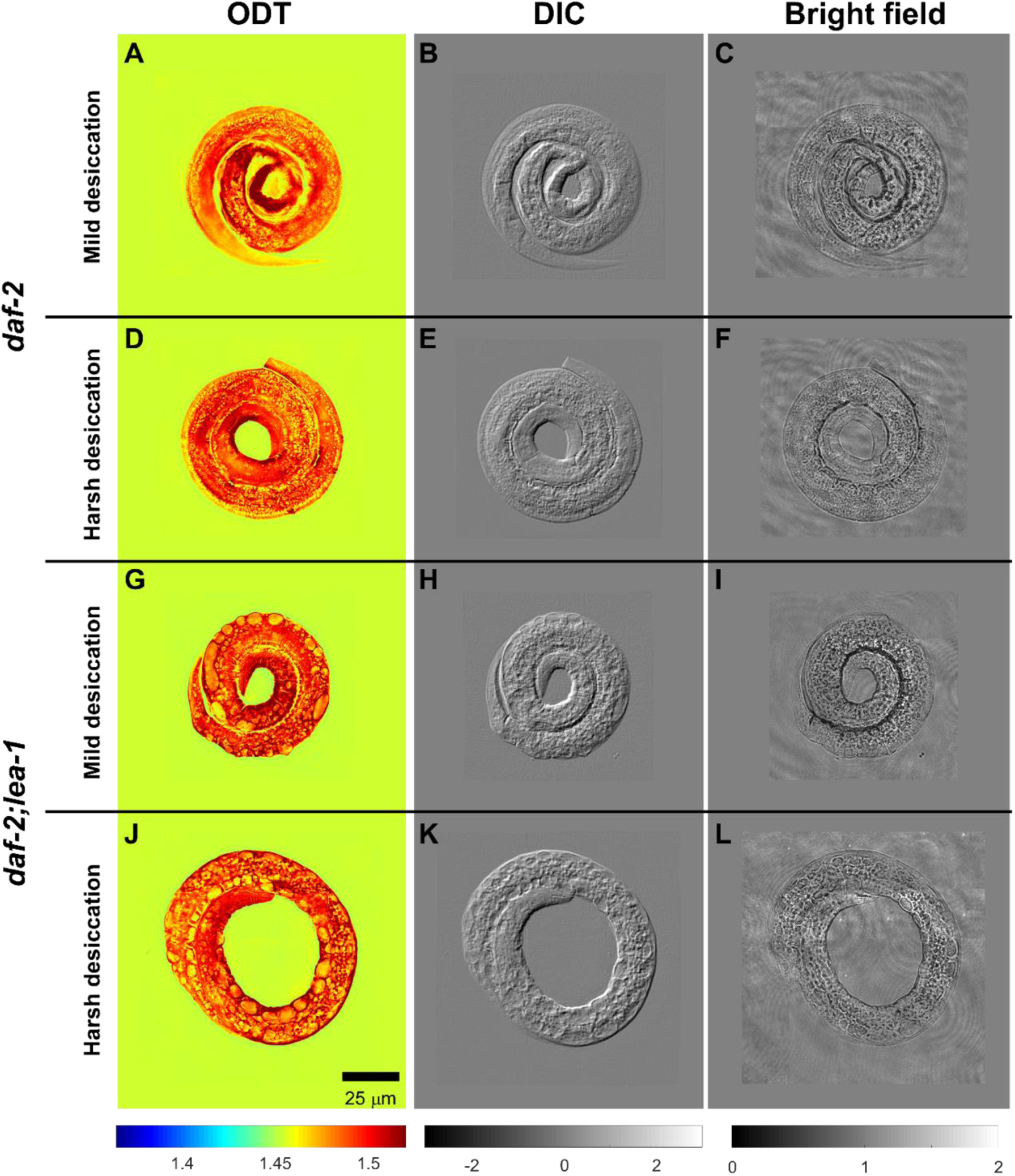
Comparison of images acquired by different microscopy modalities. (A, D, G, J) Typical central cross-sectional slices through RI tomograms, (B, E, H, K) emulated differential interference contrast (DIC) images, and (C, F, I, L) bright field images of *C. elegans* dauer larvae. (A – C) represent wild type after mild (98% RH) desiccation, (D – F) wild type after harsh (60% RH) desiccation, (G – I) *lea-1* deletion mutants *daf-2;lea-1* after mild desiccation, and (J – L) *daf-2;lea-1* after harsh desiccation. Color scale shows RI. Grey scales show light intensity in the arbitrary unit.

**Supplementary Figure 4.**
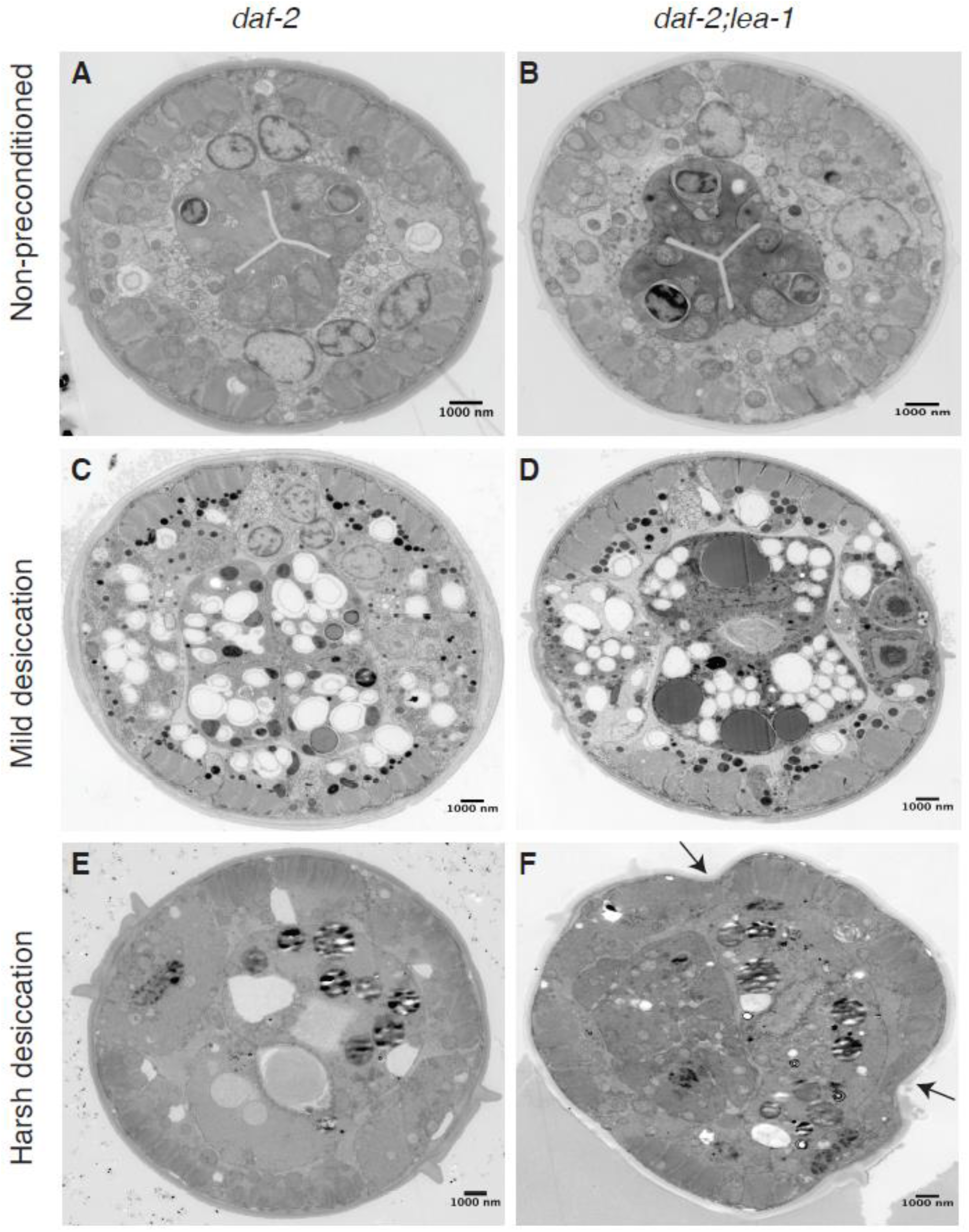
Electron micrographs of (A, C, E) wild type (*daf-2*) and (B, D, F) *lea-1* deletion mutant (*daf-2;lea-1*) in (A, B) the non-preconditioned, (C, D) mild desiccation, and (E, F) harsh desiccation conditions. The arrows in F indicate the shrinkage of the desiccated dauer larvae of *lea-1* deletion mutant.

**Supplementary Figure 5.**
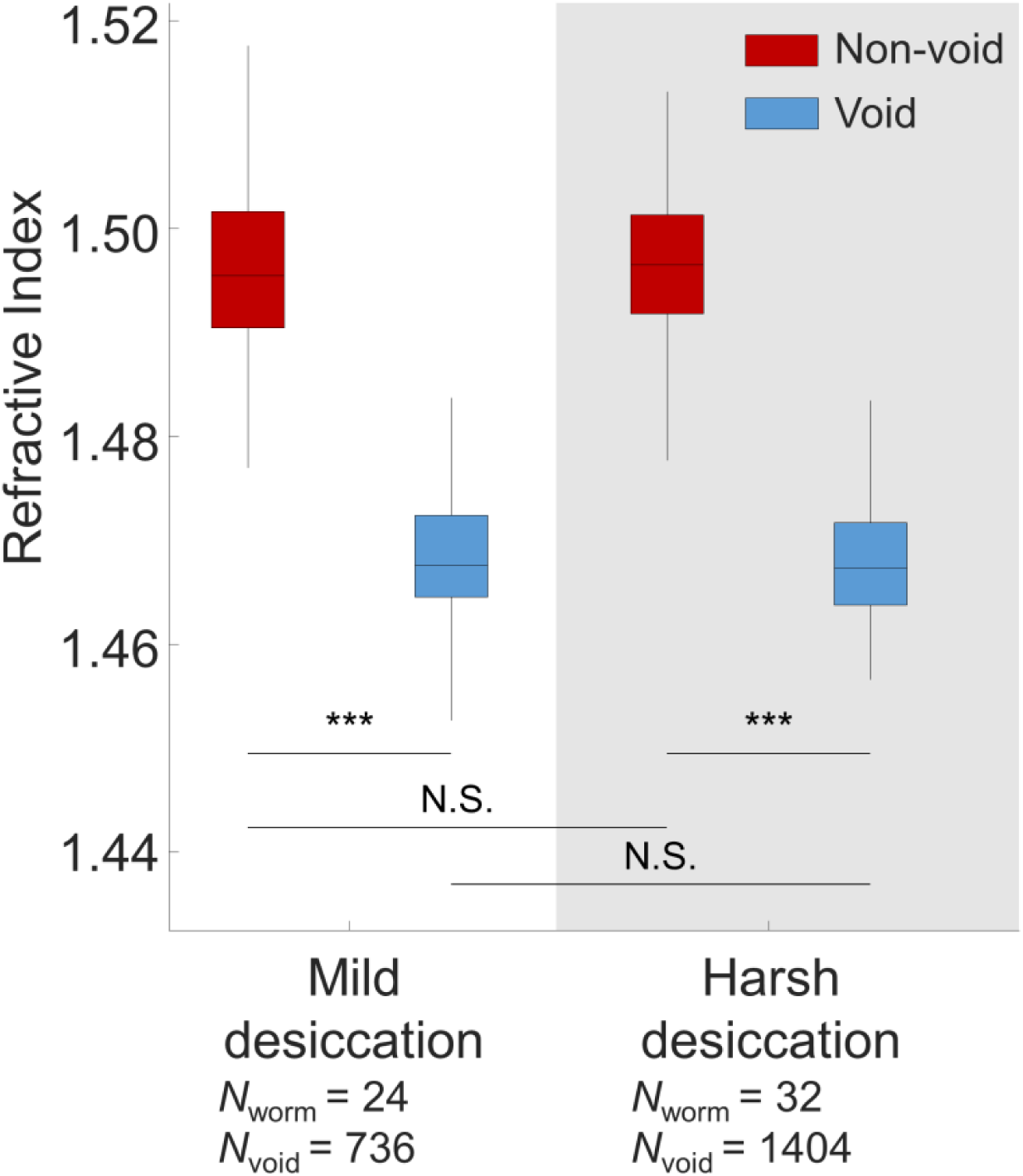
Refractive index in void regions and their periphery of *lea*-1 deletion mutant during mild and harsh desiccation conditions.

**Supplementary Figure 6.**
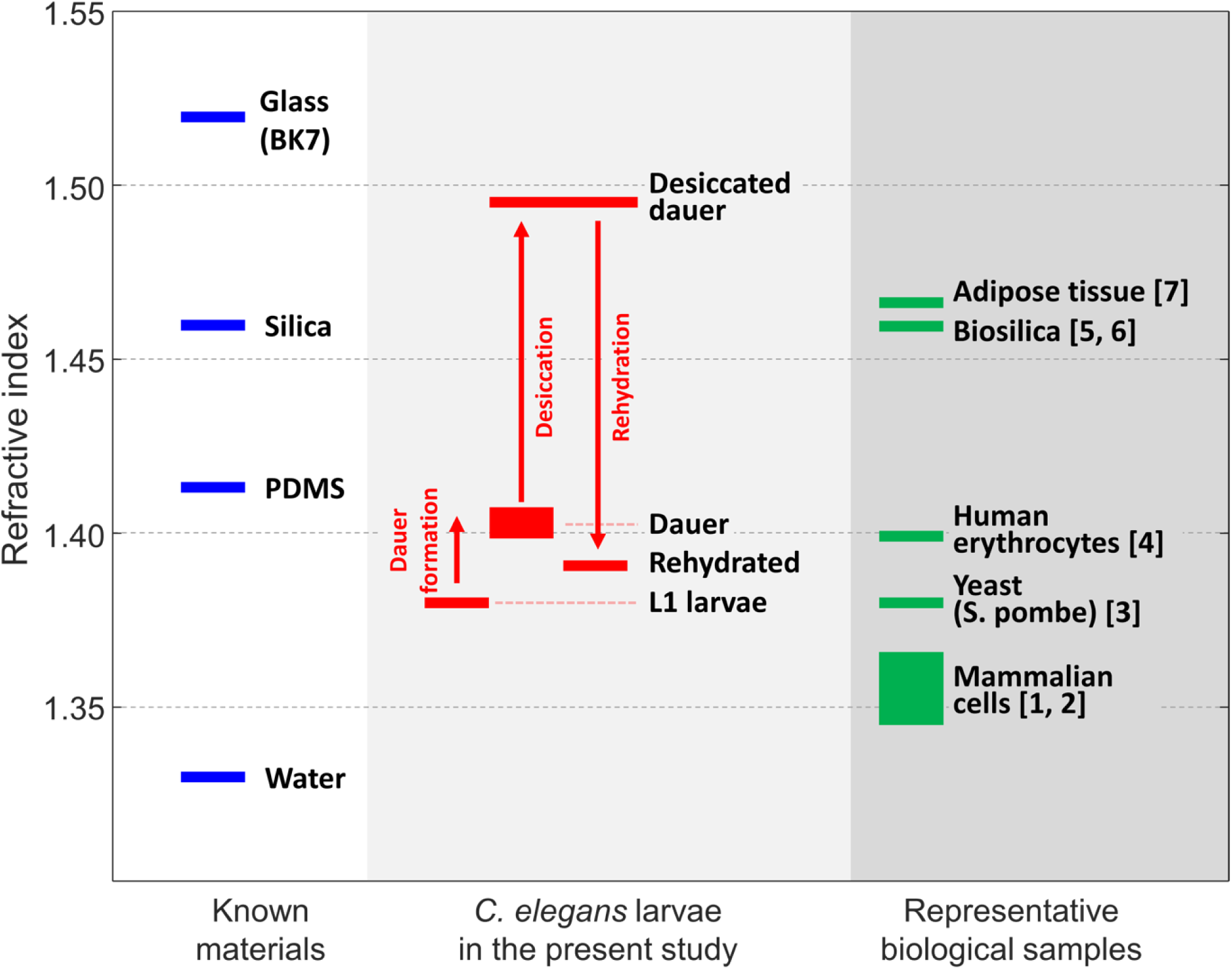
Graphical summary on the measured refractive index (RI) value of *C. elegans* larvae in the present study (center) compared to the RI of known inanimate materials (left) and representative biological samples (right).

## Supplementary Information

**Supplementary Video 1**. Visualization of the RI tomogram and rendered isosurface of a typical *C. elegans* larva at the L3 stage.

**Supplementary Video 2**. Visualization of the RI tomogram and rendered isosurface of a typical *C. elegans* dauer larva of controls after harsh desiccation (60% RH).

**Supplementary Video 3**. Visualization of the RI tomogram and rendered isosurface of a typical *C. elegans* dauer larva of trehalose deletion mutants *daf-2;ΔΔtps* after harsh desiccation (60% RH).

**Supplementary Video 4**. Visualization of the RI tomogram and rendered isosurface of a typical *C. elegans* larva of *lea-1* deletion mutants *daf-2;lea-1*after harsh desiccation (60% RH).

## Notes

https://doi.org/10.6084/m9.figshare.14483331

### Competing Interest Statement

The authors have declared no competing interest.

